# Mice lacking the cAMP effector protein POPDC1 show enhanced hippocampal synaptic plasticity

**DOI:** 10.1101/2021.05.12.443909

**Authors:** Mahesh Shivarama Shetty, Laurence Ris, Roland F. R. Schindler, Keiko Mizuno, Laura Fedele, Karl Peter Giese, Thomas Brand, Ted Abel

**Affiliations:** Department of Neuroscience and Pharmacology, Carver College of Medicine, University of Iowa, 51 Newton Road, 2-471B Bowen Science Building, Iowa City, IA 52242, United States; Iowa Neuroscience Institute, Carver College of Medicine, University of Iowa, 169 Newton Road, 2312 PBDB, Iowa City, IA 52242, United States; Department for Neuroscience, University of Mons, Research Institute for Health Sciences and Technology, 7000 Mons, Belgium; National Heart and Lung Institute, Imperial College London, London W12 ONN, UK; Department for Neuroscience, King’s College, London SE5 9NU, UK

**Keywords:** cAMP signaling, BVES, LTP, memory, phosphodiesterases

## Abstract

Extensive research has uncovered diverse forms of synaptic plasticity and a wide array of molecular signaling mechanisms that act as positive or negative regulators. Specifically, cAMP-dependent signaling pathways have been crucially implicated in long-lasting synaptic plasticity. In this study, we examine the role of POPDC1 (or BVES), a cAMP effector protein expressed in brain, in modulating hippocampal synaptic plasticity. Unlike other cAMP effectors, such as PKA and EPAC, POPDC1 is membrane-bound and the sequence of the cAMP-binding cassette differs from canonical cAMP-binding domains. These properties suggest that POPDC1 may have a unique role in cAMP-mediated signaling underlying synaptic plasticity. Our results show that POPDC1 is enriched in hippocampal synaptoneurosomes. Acute hippocampal slices from *Popdc1* knockout (KO) mice exhibit enhanced long-term potentiation (LTP) induced by a variety of stimulation paradigms, particularly in response to weak stimulation paradigms that in slices from wildtype mice induce only transient LTP. Furthermore, *Popdc1* KO mice did not display any further enhancement in forskolin-induced LTP observed following inhibition of phosphodiesterases (PDEs), suggesting a possible modulation of cAMP-PDE signaling by POPDC1. Taken together, these data reveal POPDC1 as a novel player in the regulation of hippocampal synaptic plasticity and as a potential target for cognitive enhancement strategies.

## Introduction

Understanding the molecular mechanisms involved in learning and memory represents a significant challenge in neuroscience. Studies over the years have identified activity-dependent synaptic plasticity as one of the key cellular mechanism of memory (Neves et al. 2008). Extensive research, both in invertebrate and vertebrate species has uncovered diverse forms of synaptic plasticity and the underlying complex molecular signaling pathways. Interestingly, this has revealed the existence of both positive and negative regulators of synaptic plasticity and memory (Abel et al.1998).

The cyclic 3’,5’-cyclic adenosine monophosphate (cAMP)-dependent signaling pathway plays a crucial role in persistent forms of synaptic plasticity (Abel et al. 1997; Frey et al. 1993). cAMP signaling is involved in a wide variety of cellular processes and subject to extensive spatiotemporal control to achieve signal- and cell type-specific responses. Subcellular compartment-specific nanodomain signaling complexes are formed in cells with the help of multiple A-kinase anchor proteins (AKAPs) (Bauman et al. 2004), which bind cAMP effector proteins such as protein kinase A (PKA) or exchange factor directly activated by cAMP (EPAC), adenylyl cyclases, phosphodiesterases (PDEs) and target proteins (Houslay 2010; Schleicher and Zaccolo 2020; Zaccolo et al. 2021). The concerted actions of these proteins lead to the transient activation of downstream molecular targets such as the α-amino-3-hydroxy-5-methyl-4-isoxazolepropionic acid (AMPA) receptor or the voltage-gated calcium channel Ca_V_1.2 in response to specific patterns of synaptic activity (Patriarchi et al. 2018).

The Popeye domain-containing (POPDC) proteins are a novel class of cAMP effector proteins characterized by an evolutionarily conserved Popeye domain (PF04831) that functions as a high-affinity cAMP binding domain (Brand 2018; Schindler and Brand 2016; Swan et al. 2019). The family comprises three members: POPDC1 (also known as BVES), POPDC2 and POPDC3 (Andree et al. 2000). POPDC1 is the most studied protein among the isoforms. The POPDC proteins display overlapping but isoform-specific expression patterns (Froese et al. 2012). Although POPDC genes are abundantly expressed in heart and skeletal muscle, where they have been primarily studied, they are also found in the central and autonomous nervous system (Andree et al. 2000; Hager and Bader 2009; Vasavada et al. 2004). POPDC proteins have a short extracellular domain followed by three transmembrane domains (Brand 2018; Schindler and Brand, 2016; Swan et al. 2019). The carboxyterminal domain of POPDC proteins is isoform-specific, variable in length, contains regions of low complexity and is predicted to be partly disordered (Froese et al. 2012). Constitutive knockouts of *Popdc1* and *Popdc2* have been engineered in mice and cardiac arrhythmia (stress-induced sinus node bradycardia) and impaired muscle regeneration have been reported in these mutants (Andrée et al. 2002; Froese et al. 2012). Likewise, patients carrying mutations in *POPDC1, POPDC2* or *POPDC3* develop cardiac arrhythmia (AV-block) and/or muscular dystrophy phenotypes (De Ridder et al. 2019; Indrawati et al. 2020; Rinne et al. 2020; Schindler et al. 2016; Vissing et al. 2019).

While expression of POPDC1 in the brain has been reported previously (Andree et al. 2000; Vasavada et al. 2004; Hager and Bader 2009), until now, no study has specifically looked into the expression pattern or determined the function of POPDC1 in the nervous system. Given the diverse roles of the cAMP signaling in the nervous system, we predict that POPDC1 might have specific functions in the modulation of cAMP signaling events in neurons. The fact that the phosphate-binding cassette of the Popeye domain differs from canonical ones further suggests a unique role, which differs from that of other cAMP effectors. Additionally, some of the POPDC1-interacting proteins identified thus far, such as the two-pore potassium channel TREK-1 (Froese et al. 2012), the synaptobrevins VAMP2 and 3 (Hager et al. 2010), zonula occludens-1 (ZO1) (Osler et al. 2005), dystrophin (Schindler et al. 2016) and N-myc downstream-regulated gene 4 (NDRG4) (Benesh et al. 2013), have all been implicated in neuronal function. This suggests that in addition to its role in striated muscle, POPDC1 might also have specific functions in neuronal tissue. Here, we investigate the role of POPDC1 in hippocampal synaptic plasticity using constitutive *Popdc1* KO mice. We show that POPDC1 is prominently expressed in the hippocampus and preferentially enriched in the synaptoneurosomal fraction, suggesting its synaptic localization. Our results show that *Popdc1* KO mice display enhancements in specific forms of synaptic plasticity. These findings provide the first evidence for an important role of the cAMP effector protein POPDC1 in hippocampal function.

## Materials and Methods

All animal procedures were carried out in accordance with National Institutes of Health regulations for the care and use of animals in research and were approved by the Institutional Animal Care and Use Committees at the University of Iowa, Imperial College London, University of Mons, and King’s College London.

### Popdc1 knockout mice

The generation of the mouse *Popdc1* KO mutant has been described previously (Andrée et al. 2002). Briefly the coding sequence of *Popdc1* was substituted by a lacZ reporter gene carrying a nuclear localization signal (NLS) through homologous recombination. The *Popdc1* KO mutation is kept on a C57BL/6J background.

### X-gal staining

Brains were harvested from heterozygous *Popdc1* KO mice and fixed with 2% (v/v) formaldehyde, 0.2% (v/v) glutaraldehyde, 0.02% (v/v) Nonidet P[box]40 (NP[box]40), 0.01% (v/v) sodium desoxycholate in PBS for 4h at 4°C. The fixed brains were washed three times with PBS and then incubated at 4°C with 20% and 30% sucrose. The brains were mounted on a cryomold with OCT compound and stored at −80°C until sectioning. 20-40 µm cryosections were mounted on glass slides (Superfrost plus). Sections were incubated with staining buffer (2 mM MgCl_2_, 0.01% (v/v) sodium desoxycholate, 0.02% NP40) and then stained with 5-bromo-4-chloro-3-indolyl-b-D-galactopyranoside (X-Gal) at 1 mg/ml in staining buffer containing 5 mM K_3_[Fe(CN)_6_] and 5 mM K_4_[Fe(CN)_6_] at 37°C until color has developed. Sections were counterstained with 0.1% Nuclear Fast Red in 5% aluminum sulfate, dehydrated through an ascending series of ethanol, immersed in xylene, and coverslipped with Entellan (E. Merck, Switzerland).

### Synaptosomal purification and Western blot analysis

All steps of synaptosomal preparation were performed as described (Engmann et al. 2010). Briefly, hippocampi from 6 weeks old wildtype mice were isolated and homogenized in 0.3 M sucrose, 40 mM Tris, pH 7.4, phosphatase and protease inhibitors (1/100 Phosphatase Inhibitor Cocktail 2 (Sigma), 250 nM okadaic acid, 100 nM fenvalerate, Protease Inhibitor Pill complete EDTA-free (Roche)), followed by two low-speed centrifugations at 1400 x g for 10 min. The pooled supernatants were spun down at high speed at 13800 x g for 10 min and the pellet (P2) was purified on a sucrose gradient (0.8 M/1.0 M/1.2 M in 40 mM Tris, pH 7.4, protease- and phosphatase inhibitors; 2 h at 82500 x g in a SW41 rotor at 4 °C). The synaptosomal (interface between 1.0 and 1.2 M sucrose) and cytoplasmic fractions were collected and diluted 3-fold with water. Two independent synaptosomal preparations were utilized. Protein concentration of each sample was determined by Bradford and subjected to gel electrophoresis and Western blot analysis. Proteins were size separated and transferred onto nitrocellulose membrane (BioTrace NT, Pall). The membrane was washed with TBST (50 mM Tris-HCl, pH 7.4, 150 mM NaCl, 0.1% [v/v] Tween-20) and blocked in 5% (w/v) low-fat milk in TBST for 1 hour at room temperature and subsequently incubated with POPDC1 antibody (sc-49889, Santa Cruz Biotechnology Inc.) overnight at 4°C. After several washes, the blots were incubated for 1 hour at room temperature with horseradish peroxidase–coupled anti-goat (PI-9500, Vector Laboratories) antibody. After washing, signals were detected using a Chemidoc gel imaging system (BioRad).

### Electrophysiology

Homozygous *Popdc1* mutants and littermate wildtype mice (both male and female) of 2-5 months of age were used. Mice were euthanized by cervical dislocation and the brain was quickly dissected into cold artificial cerebrospinal fluid (aCSF) (124 mM NaCl, 4.4 mM KCl, 1 mM NaH_2_PO_4_, 2.5 mM CaCl_2_.2H_2_O, 1.3 mM MgSO_4_.7H_2_O, 26.2 mM NaHCO_3_ and 10 mM D-glucose; pH ∼7.4) being continuously bubbled with carbogen (95% O_2_, 5% CO_2_). Isolation of hippocampi and preparation of acute hippocampal slices was performed as described (Shetty et al. 2015). Transverse acute hippocampal slices of 400-µm thickness were prepared from both hippocampi using a manual McIlwain slicer (Stoelting, Wooddale, IL, USA). The slices were quickly transferred onto a net insert in an interface recording chamber (Fine Science Tools, Foster City, CA) and incubated at 28°C in a humidified carbogen atmosphere for at least 2-3 hours before starting the recordings. The slices were perfused at 1 mL/min with oxygenated aCSF throughout the experiments. Field excitatory post-synaptic potentials (fEPSPs) were recorded in CA1 stratum radiatum by stimulating Schaffer collaterals with a monopolar, lacquer coated stainless-steel electrode (5 MΩ resistance, A-M Systems) and recording with an aCSF-filled glass microelectrode (2–5 MΩ resistance). Test stimulation was a biphasic, constant current pulse (100 µs duration) delivered every minute at a stimulation intensity that evoked ∼40% (or ∼50% in LTD experiments) of the maximal fEPSP amplitude as determined by an input-output curve (stimulation intensity v/s fEPSP amplitude) in each experiment. In all experiments, a stable baseline was recorded for at least 20 minutes before LTP induction or drug application. Paired-pulse facilitation (PPF) was examined at various interpulse intervals (300–25 ms). In all the electrophysiological data, ‘*n’* represents the number of mice, except in PKA inhibitor experiments where the ‘*n*’ represents the number of slices.

### Synaptic plasticity paradigms

1-train LTP stimulation comprised of a single 100 Hz train (1 s duration; pulse width 1 ms). Spaced 4-train LTP paradigm comprised of four 100 Hz, 1 s trains (pulse width 1 ms) delivered with an intertrain interval of 5 minutes. Weak theta-burst LTP stimulation (TBS) consisted of 2 bursts delivered at 5 Hz with each burst containing 4 pulses delivered at 100 Hz (pulse width 1 ms). For long-term depression (LTD) experiments, 2-3 months old mice (both male and female) were used. LTD was induced by 1 Hz stimulation for 15 minutes (900 pulses; pulse width 0.2 ms). All stimulation protocols were delivered at the baseline stimulation intensity. Forskolin (Sigma-Aldrich; #F3917) was prepared as a 50 mM stock in DMSO and bath applied at a final concentration of 50 µM in aCSF for 15 minutes (final DMSO concentration of 0.1%). The phosphodiesterase inhibitor 3-Isobutyl-1-methylxanthine (IBMX) (Sigma-Aldrich; #I5879) was prepared as a 90 mM stock in DMSO and bath applied at a final concentration of 30 µM in aCSF for 15 minutes. The PKA inhibitor KT5720 (Tocris; #1288) was prepared as a 1 mM stock in DMSO and bath applied at a final concentration of 1 µM in aCSF for 30 minutes (final DMSO concentration of 0.1%). As a vehicle control, 0.1% DMSO in aCSF was used.

### Electrophysiology Data Analysis

Data was acquired using Clampex 10 and Axon Digidata 1440 digitizer (Molecular Devices, Union City, CA) at 20 kHz and were low-pass filtered at 2 kHz with a four-pole Bessel filter. Data analysis was performed using Clampfit 10. Data plots and statistical analysis was performed using GraphPad Prism 9. For each slice, the fEPSP slopes were normalized against the average slope over the 20 minutes baseline. Data from replicate slices from the same mouse were averaged when comparing between genotypes. In electrophysiology experiments, ‘n’ represents the number of mice when the comparison is made between genotypes whereas for comparing two conditions within the same genotype, the ‘n’ represents slices. Data distribution was checked by normality tests. The maintenance of LTP was compared between two groups using independent t-tests on the last 20-min of the recordings. Input-Output and PPF data were analyzed using two-way repeated-measures ANOVA with Sidak’s test for multiple comparisons. Data are presented as mean ± SEM. Differences were considered statistically significant when Alpha (P) < 0.05.

### cAMP ELISA assay

*Popdc1* KO and WT littermate mice of 2-3 months age (both male and female) were euthanized by cervical dislocation and the hippocampal slices were prepared as described for the electrophysiology experiments. Slices from each mouse were incubated in two independently perfusable wells of the same interface incubation chamber (same chambers used for the electrophysiology experiments). Following 2 hours of recovery incubation in oxygenated aCSF, forskolin (50 µM) or vehicle (0.1% DMSO) dissolved in aCSF was bath applied to the slices for 15 minutes. Ten minutes after the end of the drug application, slices were quickly collected in a pre-chilled Eppendorf tube on dry ice, flash frozen in liquid nitrogen and stored at −80°C. On the day of the assay, samples were homogenized in 0.1M HCl using stainless steel beads. The homogenate was centrifuged at 600 x g for 10 minutes and the supernatant was immediately processed in triplicates for cAMP ELISA using the direct cAMP ELISA kit for intracellular cAMP quantification (#ADI-900-066; Enzo Life Sciences, Farmingdale, NY, USA) following manufacturer’s instructions. The cAMP levels were normalized to the total protein concentration in the supernatant determined by Bradford Assay using BSA standard curve (Bio-Rad). The final cAMP concentration is expressed as pmol cAMP per mg of total protein. The vehicle-treated slices are denoted as ‘Basal’ group and the forskolin-treated slices are denoted as ‘FSK’ group. The differences between the basal and FSK groups in WT and *Popdc1* KO mice were analyzed by two-way ANOVA followed by Šídák’s multiple comparisons tests.

## Results

### POPDC1 is expressed in the hippocampus and is enriched in synaptoneurosomes

The expression of *Popdc1* in the hippocampus was investigated by staining for beta galactosidase using coronal brain sections of heterozygous *Popdc1* KO mice, which carry a nuclear localization signal (NLS)-LacZ-reporter gene in the *Popdc1* locus (Andrée et al. 2002). A strong expression of *Popdc1* in the CA1, CA2, CA3 and dentate gyrus (DG) regions was observed (**Fig. 1A**). Expression of *Popdc1* was also seen in other parts of the brain (data not shown). We then assessed the subcellular localization of POPDC1 in the hippocampus and found that it was preferentially enriched in the synaptoneurosomal fraction. A prominent immunoreactive band at approximately 75 kDa was present in synaptoneurosomal fraction but not in the cytosolic fraction (**Fig. 1B**). The higher molecular weight of the immunoreactive band in brain tissue compared to the heart is probably due to tissue-specific glycosylation as previously suggested (Vasavada et al. 2004). This suggests that POPDC1 localizes preferentially near the synapses indicating its potential role in synaptic function.

**Figure 1.**
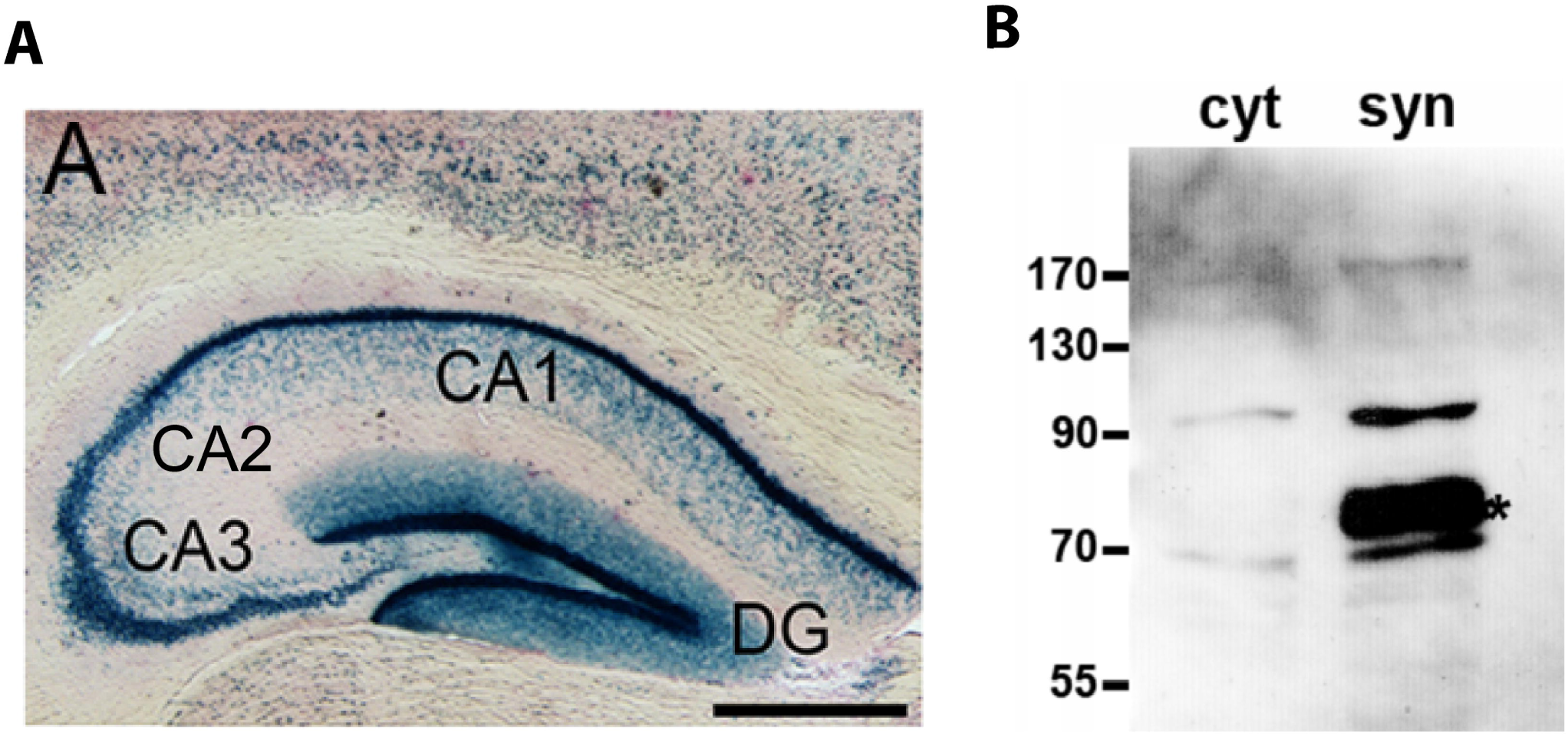
*Popdc1* is expressed in all the hippocampal subregions and the protein preferentially localizes near the synapses. **(A)** β-galactosidase staining of a coronal section of the brain of a heterozygous *Popdc1* KO mouse. Only the hippocampal region is depicted. (**B**) Western blot analysis of cytoplasmic fraction (cyt) and synaptoneurosomes (syn) isolated from the wildtype mouse hippocampus. Asterisk demarcates the immunoreactive band for POPDC1. CA1, CA2, CA3 – Cornu Ammonis areas 1, 2 and 3; DG - dentate gyrus. Scale bar in (A): 0.5 mm.

### *Popdc1* KO mice show PKA-dependent enhancement in LTP in response to submaximal high frequency stimulation

We next examined the impact of the loss of POPDC1 on long-term potentiation (LTP) at the hippocampal CA1 Schaffer collateral synapses using acute slices. The Schaffer collateral fibers were stimulated and the field-EPSP responses were recorded in the CA1 stratum radiatum. In slices from WT animals, stimulation with a single train of 100 Hz, 1 s protocol elicited a transient potentiation (1-train LTP) **(Fig. 2A)**. It is thought that 1-train LTP is independent of both protein synthesis and PKA activation, and decays to baseline levels within 1-2 hours (Frey et al. 1993; Huang and Kandel 1994; Nguyen and Woo 2003). Interestingly, the same protocol resulted in long-lasting LTP in slices from *Popdc1* KO mice **(Fig. 2A, 2B)**. Next, we assessed LTP induced by a stronger protocol comprising four 100 Hz, 1 s trains spaced at 5 mins (spaced 4-train LTP). This stimulation paradigm induces a saturating LTP that is long-lasting and is dependent on transcription, translation and PKA activation (Abel et al. 1997; Huang and Kandel 1994; Malleret et al. 2001; Nguyen et al. 1994). The spaced 4-train LTP was similar and non-decremental in slices from both the *Popdc1* KO and WT littermates **(Fig. 2C, 2D).**

**Figure 2.**
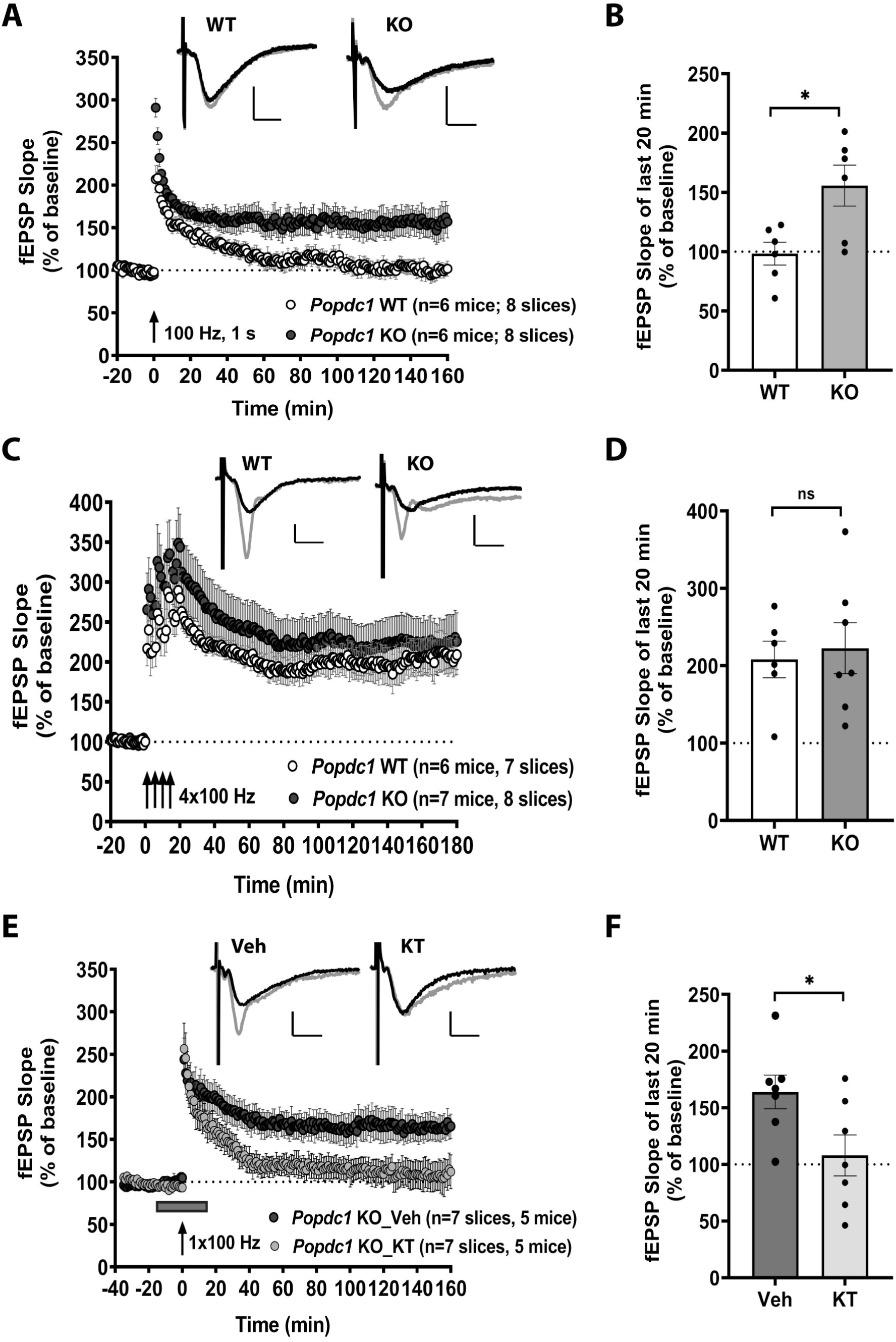
*Popdc1* KO mice show PKA-dependent long-lasting LTP in response to submaximal high frequency stimulation. **(A)** LTP induced by a single 100 Hz train (1-train LTP) is persistently enhanced in slices from *Popdc1* KO mice (n=6 mice; 4 males, 2 females) compared to WT littermates (n=6 mice; 2 males, 4 females). **(B)** The mean fEPSP slope over the last 20 min recordings was significantly higher in *Popdc1* KO (155.7±17.33%) compared to WT littermates (98.33±9.6%); Unpaired t-test, two-tailed, t=2.897, df=10, P=0.0159. **(C)** Persistent LTP induced by repeated 100 Hz stimulation trains (spaced-4-train protocol) was similar in slices from *Popdc1* KO mice (n=7 mice; 4 males, 3 females) compared to WT littermates (n=6 mice; 4 males, 2 females). **(D)** The mean fEPSP slope over the last 20 min recordings was not significantly different in the *Popdc1* KO (222.6±32.83%) compared to WT littermates (208.1±23.64%); Unpaired t-test, two-tailed, t=0.3464, df=11, P=0.7356. **(E)** The enhanced 1-train LTP observed in slices from *Popdc1* KO mice was dependent on PKA activation. Bath application of the PKA inhibitor KT5720 (1 µM) during LTP induction prevented the LTP enhancement (n=7 slices; 5 mice; 4 males, 1 female) whereas vehicle (DMSO) treatment did not (n=7 slices; 5 mice; 2 males, 3 females). **(F)** The mean fEPSP slope over the last 20 min recordings was significantly lower after KT5720 treatment (107.9±18.07%) compared to slices receiving only vehicle (163.9±14.85%); Unpaired t-test, two-tailed, t=2.396, df=12, P=0.0338. In **A, C** and **E,** representative fEPSP traces during the baseline (black traces) and at the end of the recording period (gray traces) for each group are shown. Calibration bars for all the traces: 2 mV vertical; 5 ms horizontal.

Because PKA activation is crucially involved in long-lasting forms of LTP (Abel et al. 1997; Nguyen and Woo 2003), we then investigated the PKA dependence of the persistent 1-train LTP seen in the *Popdc1* KO mice. Bath application of the PKA inhibitor KT5720 to the slices during the 1-train stimulation (15 min before and 15 min after) blocked the enhancement in LTP compared to vehicle-treated control slices **(Fig. 2E, 2F).** This suggests that the enhanced LTP seen in the *Popdc1* KO mice requires PKA activity.

### Basal synaptic transmission and short-term plasticity are not altered in *Popdc1* knockout mice

Next, we examined whether the loss of *Popdc1* alters basal synaptic transmission in hippocampal neurons by comparing input-output responses. The stimulus-response curves evoked with progressively increasing stimulus intensities (5 μA - 70 μA) showed no significant differences both in the field-EPSP amplitudes **(Fig. 3A)** and in presynaptic fiber volley amplitudes **(Fig. 3B)** between the wildtypes and knockouts. We also investigated paired-pulse facilitation, a form of short-term synaptic plasticity and an index of presynaptic activity and release probability where two stimuli given in close succession lead to the facilitation in the response to the second stimuli (Jackman and Regehr 2017; Regehr 2012; Zucker and Regehr 2002). At all the interstimulus intervals tested (ranging from 300 ms to 25 ms), no significant differences in the facilitation, calculated as paired-pulse ratio (PPR= Amplitude of fEPSP2/Amplitude of fEPSP1), was observed between slices from WT and *Popdc1* KO animals **(Fig. 3B)**. Together, these results suggest that POPDC1 may not have a significant role in the maintenance of basal synaptic transmission and very short-term plasticity.

**Figure 3.**
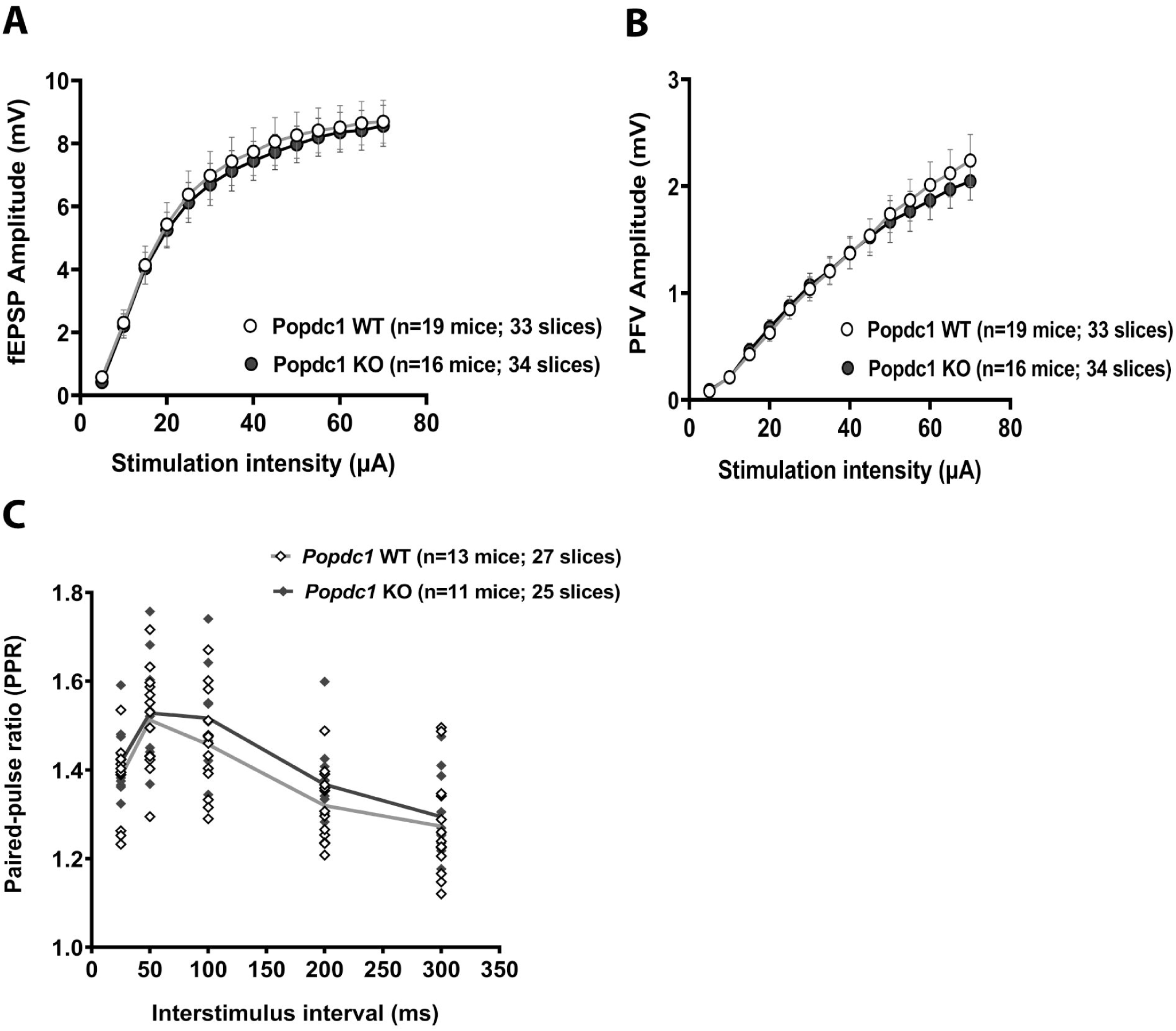
*Popdc1* KO mice show no alterations in basal synaptic transmission and paired-pulse facilitation. **(A)** Input-output relation plot of the fEPSP amplitude versus the incremental stimulation intensity in *Popdc1* KO mice (n=16 mice; 6 females, 10 males) and WT littermates (n=19 mice; 9 females; 10 males). Two-way repeated measures ANOVA showed no significant differences between the genotypes F (1,33) = 0.064; P=0.801. Multiple comparisons using Sidak’s test showed no significant differences between groups at any of the stimulation intensities tested (P>0.05). **(B)** Input-output relation plot of the presynaptic fiber volley (PFV) amplitude versus the incremental stimulation intensity in *Popdc1* KO mice (n=16 mice; 6 females, 10 males) and WT littermates (n=19 mice; 9 females; 10 males). Two-way repeated measures ANOVA showed no significant differences between the genotypes F (1,33) = 0.044; P=0.835. Multiple comparisons using Sidak’s test showed no significant differences between the groups at any of the stimulation intensities tested (P>0.05). **(C)** Paired-pulse facilitation (calculated as Paired-pulse ratio (PPR) = amplitude of fEPSP2/amplitude of fEPSP1) measured over a range of interpulse intervals (25 ms - 300 ms) in *Popdc1* KO mice (n=11 mice; 5 females, 6 males) and WT littermates (n=13 mice; 7 females; 6 males). Two-way repeated measures ANOVA showed no significant differences between the genotypes F (1,22) = 1.061; P=0.3141. Multiple comparisons using Sidak’s test showed no significant differences between the groups at any of the interpulse intervals tested (P>0.05).

### *Popdc1* KO mice show enhancement in potentiation induced by forskolin but not with simultaneous inhibition of phosphodiesterases

Because we observed a PKA-dependent enhancement in 1-train LTP in the *Popdc1* K*O* mice, we next examined the long-lasting potentiation induced by raising cAMP levels using forskolin (FSK), a direct pharmacological activator of adenylyl cyclase (Chavez-Noriega and Stevens 1992; Huang and Kandel 1994; Otmakhov et al. 2004). Bath application of FSK (50 μM) resulted in significantly enhanced potentiation in slices from *Popdc1* KO animals compared to WT **(Fig. 4A and 4B).** Interestingly however, the potentiation induced by the co-application of FSK (50 μM) and the pan-phosphodiesterase inhibitor IBMX (30 μM) showed no further enhancement and was similar in the WT and *Popdc1* KO slices **(Fig. 4C and 4D)**. Because POPDC1 is a cAMP effector protein and we observed enhanced FSK-induced potentiation in the *Popdc1* KO slices, we tested if the loss of POPDC1 leads to enhanced cAMP levels. We used a cAMP ELISA assay to measure gross intracellular cAMP levels in the hippocampal slices from WT and *Popdc1* KO mice following stimulation with FSK (50 μM) or vehicle (DMSO) **(Fig. 4E)**. The results showed no statistically significant difference in the cAMP levels, either at basal conditions or following forskolin stimulation, between the slices from WT and *Popdc1* KO mice.

**Figure 4.**
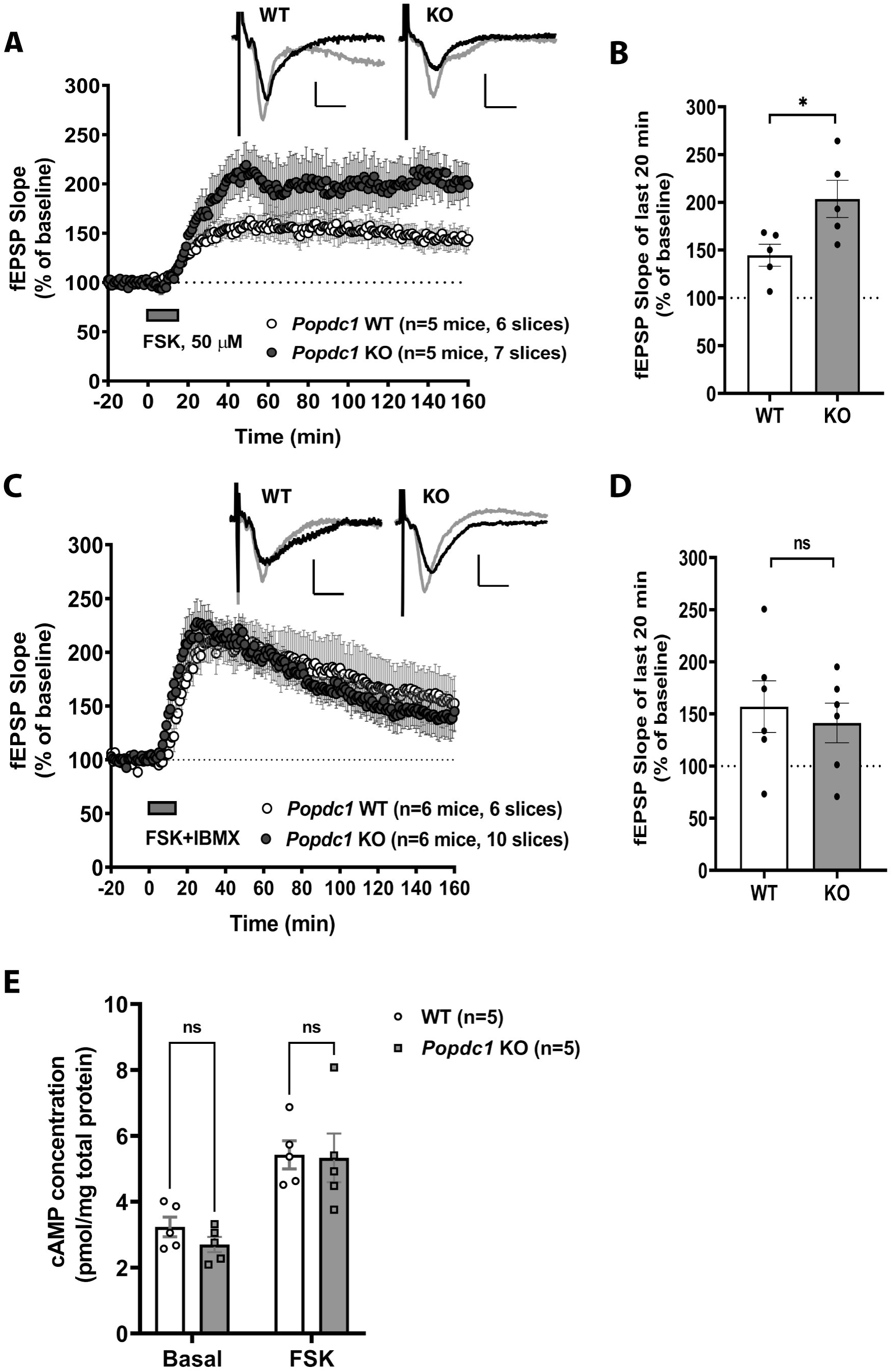
*Popdc1* KO mice show enhancement in potentiation induced by forskolin but not with simultaneous inhibition of phosphodiesterases. **(A)** Persistent potentiation induced by bath application of 50 µM forskolin (FSK) was enhanced in slices from *Popdc1* KO mice (n=5 mice; 3 males, 2 females) compared to WT littermates (n=5 mice; 2 males, 3 females). **(B)** The mean fEPSP slope over the last 20 min recordings was significantly higher in *Popdc1* KO (203.6±19.5%) compared to WT (144.6±11.5%); Unpaired t-test, two-tailed, t=2.611, df=8, P=0.0311. **(C)** Potentiation induced by co-application of 50 µM forskolin and 30 µM IBMX (FSK+IBMX) in slices from *Popdc1* KO mice (n=6 mice; 5 males, 1 female) was indistinguishable from WT littermates (n=6 mice; 5 males, 1 female). **(D)** The mean fEPSP slope over the last 20 min recordings was not significantly different in *Popdc1* KO (141.3±19.1%) compared to WT littermates (157±24.7%); Unpaired t-test, two-tailed, t=0.5028, df=10, P=0.626. In **A** and **C,** representative fEPSP traces during baseline (black traces) and at the end of the recording period (gray traces) for each group are shown. Calibration bars for all traces: 2 mV vertical; 5 ms horizontal. **(E)** The total levels of cAMP measured in hippocampal slices of WT and *Popdc1* KO mice treated with 50 μM FSK or DMSO as vehicle (Basal). Similar basal cAMP levels in *Popdc1* KO slices (2.702±0.23 pmol/mg of total protein; n=5 mice; 1 male, 4 females) compared to WT (3.238±0.30 pmol/mg of total protein; n=5 mice; 2 males, 3 females). Comparable cAMP levels in *Popdc1* KO slices (5.331±0.74 pmol/mg of total protein) and WT (5.426±0.43 pmol/mg of total protein) after FSK-stimulation; 2-way ANOVA, F(1,16) = 0.456, P=0.509; Šídák’s multiple comparisons tests between genotypes at basal or after FSK, P>0.05.

These results indicate that POPDC1 plays a role in regulating local cAMP signaling possibly by regulating the function of phosphodiesterases. Because changes in cAMP levels might be predominantly happening in localized domains near the synapses, estimation of gross cAMP levels may not have reliably captured these changes.

### Low-frequency stimulation-induced LTP is enhanced but LTD is not altered in *Popdc1* knockout mice

We further tested synaptic plasticity induced by theta-burst stimulation (TBS), a paradigm that has been proposed to resemble the rhythmic hippocampal electrical activity observed during spatial exploration (Buzsaki 2005). Because we observed an enhancement in 1-train LTP in the *Popdc1* KO mice, we used a weaker version of TBS containing only two bursts delivered at 5 Hz with each burst containing four 100 Hz pulses. This weak 2-burst TBS protocol has been shown to induce a transient potentiation whereas protocols containing more than five bursts result in persistent LTP (Kramar et al. 2009). Consistent with this, weak-TBS stimulation elicited a transient potentiation in slices from WT animals. In slices from *Popdc1* KO mice, the weak-TBS stimulation resulted in enhanced potentiation compared to slices from WT **(Fig. 5A and 5B).** On the other hand, LTD induced with a prolonged low-frequency stimulation protocol (Dudek and Bear, 1992) showed no alterations between WT and *Popdc1* KO mice **(Fig. 5C and 5D)**. Although we studied LTD in 2-3 months old mice, under our stimulation parameters and experimental conditions involving long incubation periods and aCSF composition with Ca^2+^/Mg^2+^ ratio >1, we could reliably induce LTD in slices from these mice using the 1 Hz, 15 min (900 pulses) protocol, as noted in previous studies (Billard 2010; Delgado et al. 2018; Dudek and Bear, 1992; Heynen et al. 1996; Staubli and Ji, 1996).

**Figure 5.**
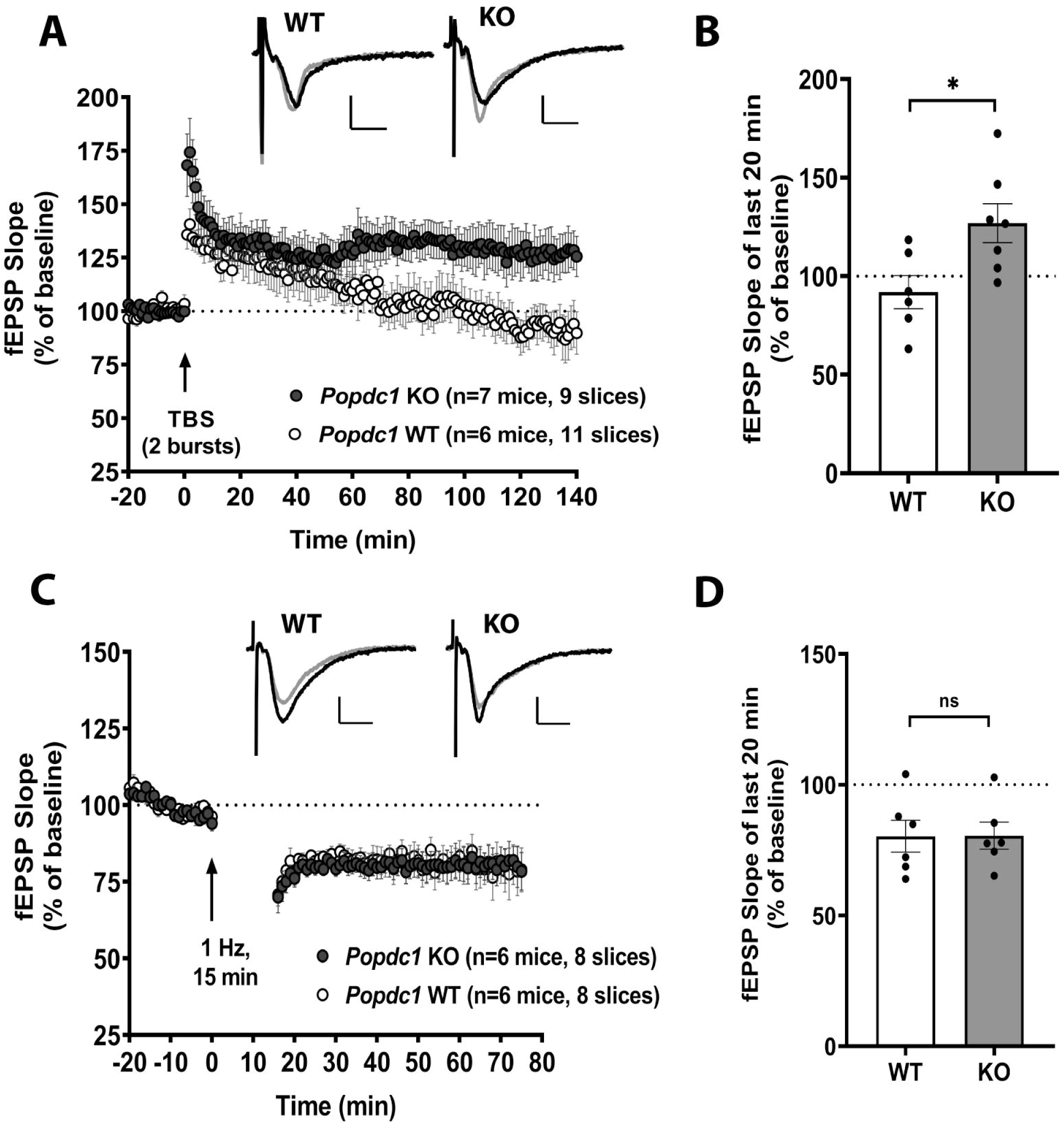
*Popdc1* KO mice show enhanced theta-burst LTP but show no alteration in LTD induced by low-frequency stimulation. **(A)** LTP induced by a weak theta-burst stimulation is enhanced in slices from *Popdc1* KO mice (n=7 mice; 5 males, 2 females) compared to WT littermates (n=6 mice; 5 males, 1 female). **(B)** The mean fEPSP slope over the last 20 min recordings was significantly higher in *Popdc1* KO (126.9±9.9%) compared to WT littermates (91.9±8.4%); Unpaired t-test, two-tailed, t=2.648, df=11, P=0.0226. **(C)** LTD induced by prolonged 1 Hz stimulation protocol was similar in slices from *Popdc1* KO mice (n=6 mice; 3 males, 3 females) compared to WT littermates (n=6 mice; 5 males, 1 female). **(D)** The mean fEPSP slope over the last 20 min recordings was not significantly different in *Popdc1* KO (80.5±5.2%) compared to WT littermates (80.3±6.1%); Unpaired t-test, two-tailed, t=0.0329, df=10, P=0.9743. In **A** and **C,** representative fEPSP traces at baseline (black traces) and at the end of the recording period (gray traces) for each group are shown. Calibration bars for all traces: 2 mV vertical; 5 ms horizontal.

## Discussion

The cAMP-mediated signaling pathway plays a crucial role in the persistent forms of synaptic plasticity (Abel and Nguyen 2008; Abel et al. 1997). Given the central role of this signaling pathway in many cellular processes, the activation of downstream targets is spatiotemporally regulated. This is achieved in a number of ways: via controlling the generation of cAMP by adenylyl cyclases, by regulating cAMP degradation through phosphodiesterases and by the subcellular targeting of cAMP effector proteins with the help of a number AKAP proteins (Zaccolo et al. 2021). The cAMP effector proteins PKA, EPAC, and hyperpolarization-activated cyclic nucleotide-gated cation channel (HCN) have been well characterized in neurons (Kelly 2018). The POPDC proteins are a recent addition to this group of cAMP effectors (Brand 2018) and until now their function in neurons has not been studied. In this study, we investigated the role of POPDC1 in synaptic plasticity in the hippocampal CA3-CA1 Schaeffer collateral synapses using mice lacking *Popdc1* (Andrée et al. 2002). Beta-galactosidase staining of the LacZ reporter gene in the *Popdc1* locus showed expression in the different subregions of hippocampus. Moreover, POPDC1 protein showed preferential enrichment in synaptoneurosomes compared to the cytosolic fraction, supporting its potential role in synaptic function.

Our results show that *Popdc1* KO mice exhibit enhanced LTP induced by a single HFS train. Similar enhancement has also been reported following pharmacological enhancement of hippocampal cAMP levels (Barad et al. 1998), in mice expressing an inhibitor of ATF4 (CREB-2) and C/EBP proteins (Chen et al. 2003) and following genetic inhibition of calcineurin (Malleret et al. 2001). The LTP enhancement was not apparent in response to strong tetanization protocols, possibly due to the saturating magnitude of potentiation induced by such paradigms or, in other words, due to a ‘ceiling effect’. Persistent LTP observed in the *Popdc1* KO following a single HFS train was found to be dependent on PKA activation. This is in line with previous studies implicating a crucial role for PKA in persistent forms of LTP where PKA activation functions like a gate between transient and persistent forms of LTP (Abel et al. 1997; Blitzer et al. 1995). We also observed similar LTP enhancement in hippocampal slices of *Popdc1* KO mice subjected to a subthreshold theta-burst stimulation (TBS) protocol. Persistent LTP induced by stronger TBS protocols (those involving more than 5 bursts) has been shown to be dependent on PKA (Nguyen and Kandel 1997) and PKA anchoring by AKAPs (Nie et al. 2007). Overall, our findings suggest that loss of *Popdc1* results in reduced LTP induction threshold. Given that aging is associated with increased LTP threshold and impairments in persistence of LTP (Bach et al. 1999), in future studies it will be interesting to investigate whether blocking POPDC1 function might be able to rescue age-related deficits.

The finding that the loss of POPDC1 results in enhancement of LTP in response to weak stimulation paradigms suggests that POPDC1 might function as a molecular brake to constrain the plasticity in the hippocampal CA1 region. Given its known role as a cAMP-effector protein, POPDC1 seems to function by regulating cAMP-PKA dependent signaling that is crucially involved in persistent forms of synaptic plasticity. Our results demonstrating no alteration of LFS-induced LTD in *Popdc1* KO mice indicate that POPDC1 is preferentially involved in the regulation of LTP and not LTD. Nevertheless, other forms of LTD induced either by other electrical or chemical stimulation paradigms need to be investigated to be conclusive about this preferential involvement.

An intriguing finding in this study is that the *Popdc1* KO mice show an enhancement in the magnitude of the long-lasting potentiation induced by FSK but no difference in potentiation is observed between the null mutant and WT when FSK was applied together with IBMX (FSK+IBMX). Both these forms of potentiation are known to be dependent on cAMP-PKA signaling (Chavez-Noriega and Stevens 1992, 1994). In slices from WT mice, FSK+IBMX application results in robust enhancement in the magnitude of potentiation compared to that induced by FSK alone. In contrast, in slices from *Popdc1* KO mice, the initial levels of potentiation are similar following FSK alone or FSK+IBMX co-application, but eventually the potentiation induced by FSK+IBMX maintains at lower levels compared to FSK alone (compare the levels of potentiation in **Fig. 4A and 4C**). This partial occlusion of potentiation with PDE inhibition indicates a link between POPDC1 and PDE function. This hypothesis is strongly supported by our recent findings showing that POPDC1 indeed directly and preferentially interacts with one of the long isoforms of PDE4, PDE4A5 (Tibbo et al. 2020), an isoform that is highly expressed in the hippocampus (McPhee et al. 2001) and implicated in hippocampal synaptic plasticity (Havekes et al. 2016; Vecsey et al. 2009; Wong et al. 2019). The interface of this interaction was mapped using peptide arrays to the Popeye domain of POPDC1 and to the UCR1 region of PDE4A (Tibbo et al. 2020). The POPDC1-PDE4A complex was suggested to be constitutively present and the interaction neither occluded the PDE4A active site nor the site of phosphorylation by PKA (Tibbo et al., 2020). Our working hypothesis, taking these findings into account, is that POPDC1 regulates local cAMP signaling near the synapses in response to synaptic activity possibly by regulating PDE4A localization and activity. Although we did not find significant differences in gross cAMP levels in the hippocampal slices of *Popdc1* KO mice either at basal activity or following FSK stimulation, we suspect that this might be due to the inability of the ELISA assay utilized here to capture local changes in cAMP nanodomains. It will be interesting to investigate the role of POPDC1 in local cAMP dynamics during synaptic activity using imaging approaches based on genetically encoded cAMP sensors (Schleicher and Zaccolo, 2018).

It is possible that the interaction of POPDC1 with other proteins also contributes to synaptic plasticity. Our work in cardiac myocytes has identified the two-pore potassium channel TREK-1 as an important mediator of POPDC1 function in the heart (Froese et al. 2012). In neurons, TREK-1 plays a crucial role in the maintenance of the resting membrane potential and thereby neuronal excitability (Enyedi and Czirjak, 2010; Honoré 2007). Studies have also implicated TREK-1 in synaptic plasticity and memory (Cai et al. 2017; Wang et al. 2020; Weng et al. 2016). However, we think that the POPDC1-TREK-1 interaction may not be important in hippocampal CA1 neurons because we observed no significant changes in basal synaptic transmission in the slices from *Popdc1* KO mice as assessed by changes in input-output relation **(Fig.3A, 3B)**. This is also supported by our results showing no changes in LFS-induced LTD in *Popdc1* KO mice, which rules out any gross changes in neuronal excitability. Nevertheless, the role of POPDC1-TREK-1 interaction in neurons needs to be investigated further using more direct approaches. It is also possible that POPDC1 might function in regulating the adenylyl cyclase activity following specific patterns of stimulation either by direct interaction or through G-proteins and it would be of interest to test this in future studies.

We have recently proposed a few working models by which POPDC1 might regulate cAMP signaling (Swan et al. 2019). In the simplest scenario (*Switch model*), POPDC proteins might form a complex with a target protein such as an ion channel and the binding status would act as a switch to alter the properties of the target protein. Support for this model comes from the interaction of POPDC1 with the potassium channel TREK1, which has been shown to be cAMP sensitive (Froese et al. 2012; Schindler and Brand 2016). Apart from its role in channel gating, POPDC1 is also involved in modulating channel trafficking and the *Cargo model* proposes that membrane transport, which is enhanced by POPDC1 in case of TREK1, may be also modulated by cAMP binding (Froese et al. 2012). Since POPDC1 is relatively abundant in many cell types and have a cAMP binding affinity similar to that of PKA, it is possible that it might also contribute to spatial control of cAMP diffusion by binding to cAMP and thereby limiting its diffusion and therefore the activation of other cAMP effector proteins. Finally, in the *Shield mode*l, POPDC proteins form complexes with interacting proteins, which include transmembrane proteins and membrane-associated proteins (Swan et al. 2019). Such an interaction might shield the interacting protein from certain post-translational modifications. The shield function of POPDC proteins could also be executed indirectly though the modulation of PDE activity, for example. Currently, we cannot definitively distinguish between these models in relation to hippocampal synaptic plasticity and it will be interesting to investigate the POPDC1 interactome near the synapse to precisely determine, how POPDC1 might modulate synaptic plasticity and memory.

In the light of our results from the forskolin and PDE inhibition experiments and from the recent findings showing direct POPDC1-PDE4A interaction (Tibbo et al. 2020), and the evidence implicating PDE4A in hippocampal synaptic plasticity (Havekes et al. 2016; Vecsey et al. 2009; Wong et al. 2019), we propose a working model for the role of POPDC1 in synaptic plasticity. In WT mice (**Fig. 6A**), POPDC1 localizes near the synapses and interacts with PDE4A long isoforms through their UCR1 region (Tibbo et al. 2020), thus acting like a PDE4A anchor to result in a spatially restricted cAMP localization. This interaction might prevent PDE4A dimerization, either directly by affecting the conformation or indirectly by sequestering the monomers, because the UCR1 region is crucial for the dimerization (Bolger et al. 2015). The monomeric PDE4s have an ‘open’ conformation and show much greater activity towards cAMP degradation compared to PDE4 dimers with a preferential ‘closed’ conformation (Cedervall et al. 2015; Perry et al. 2002). Additionally, beta-arrestins show higher affinity for open monomers of PDE4 than the closed dimers for recruiting them to the ligand-occupied beta2-adrenergic receptor, resulting in downregulation of cAMP signaling (Baillie et al. 2003; Perry et al. 2002). Overall, POPDC1-anchored PDE4A monomers near the synapse might be acting to limit cAMP elevation in response to weaker LTP-inducing stimuli, thereby preventing the activation of cAMP effectors, such as PKA. Further, in the absence of PKA activation, inhibitor-1 is kept in a dephosphorylated state by the action of calcineurin, which leads to the activation of protein phosphatase 1 (PP1). This phosphatase cascade would make sure that the LTP stimulus-induced synaptic changes, such as phosphoCaMKII-mediated AMPAR phosphorylation, are transient. In *Popdc1* KO mice (**Fig. 6B**), the absence of POPDC1 may lead to increased chances of PDE4A dimerization and loss of spatially restricted localization of PDE4A near the synapse. Now, in response to the LTP-inducing stimulus, cAMP can diffuse more widely, and the signal may last longer because PDE4 dimers have reduced catalytic activity and may not be properly localized. These changes may cause enhanced cAMP signaling leading to the activation of cAMP effectors, such as PKA. Active PKA may then phosphorylate many of its plasticity-related targets, including NMDARs, AMPARs, Inhibitor-1, leading to the activation of transcription and translation and ultimately persistent LTP.

**Figure 6.**
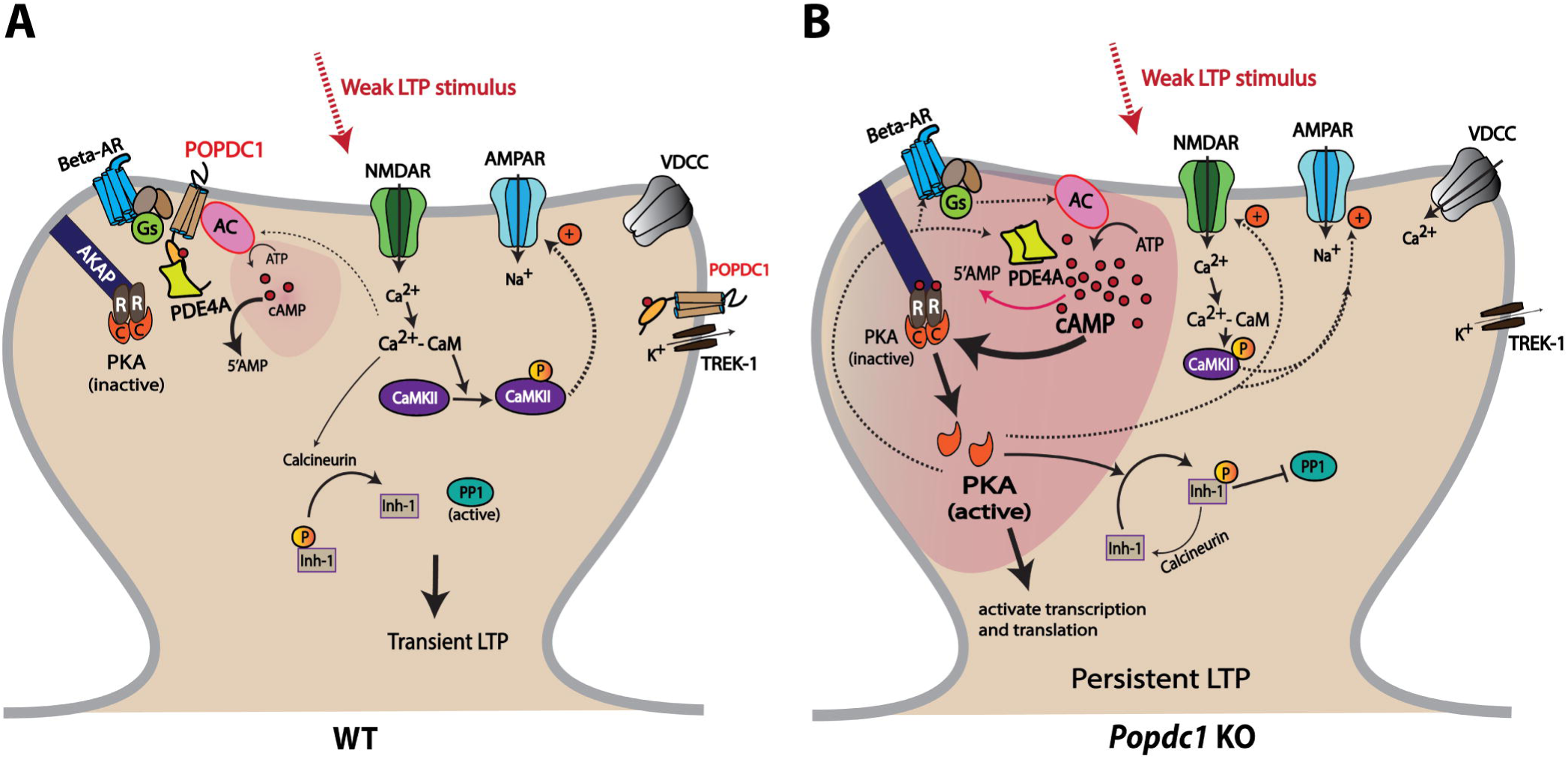
A schematic depiction of a working model to explain the enhanced LTP observed in *Popdc1* KO mice in response to submaximal high frequency stimulation. **(A)** In hippocampal neurons of WT mice, POPDC1 protein localizes near the synapse possibly interacting with other synaptic proteins involved in cAMP signaling including beta-adrenergic receptors (beta-AR), adenylyl cyclases (AC), phosphodiesterases such as PDE4A and the two-pore potassium channel TREK-1. Specifically, POPDC1 interacts with the PDE4A long isoforms through their UCR1 region, thus sequestering them in monomeric form that is highly active due to its ‘open’ conformation. A weak synaptic stimulus that induces a transient LTP leads to influx of Ca^2+^ through NMDA receptors (NMDAR) and activation of Ca^2+^-calmodulin (CaM)-dependent Kinase II (CaMKII). Active CaMKII phosphorylates AMPA receptors (AMPAR) increasing the channel conductance and also leads to increased synaptic trafficking of AMPARs. The stimulation might also induce modest increase in cAMP levels through Ca^2+^-CaM-dependent AC activation, but the cAMP is quickly degraded by the highly active PDE4 monomers, thereby preventing PKA activation. Ca^2+^-CaM-dependent activation of calcineurin keeps inhibitor-1 (Inh-1) dephosphorylated, in turn resulting in active protein phosphatase 1 (PP1). Low local cAMP levels and active phosphatase cascades only allow a short-lasting LTP. **(B)** In the hippocampal neurons of *Popdc1* KO mice, the absence of POPDC1 leads to increased PDE4A dimers that are less efficient at cAMP degradation due to the ‘closed’ conformation and also are not spatially restricted near the synapses. This results in an accumulation and persistence of local cAMP produced in response to weak LTP-inducing stimulus. This increased cAMP levels leads to an activation of PKA, which phosphorylates many of its targets such as AMPAR, NMDAR, beta-AR etc., and activates transcription and translation. This, along with inhibition of the phosphatase cascade and possibly other mechanisms involving AC inactivation, TREK-1 and voltage-dependent calcium channels (VDCC), results in persistent LTP.

Taken together, our findings identify POPDC1, a unique cAMP effector protein, as a novel player involved in the regulation of hippocampal synaptic plasticity and pave the way for future studies aimed at understanding its function in learning and memory. Although it is indeed tantalizing to investigate the role of POPDC1 in memory, some of the phenotypes displayed by the constitutive *Popdc1* knockout mice, such as the stress-induced sinus node bradycardia (Froese et al. 2012) and an overall hyperexcited state, makes this mouse line not suitable for use in behavioral memory tasks. The interaction of POPDC1 with PDE4A5 and its potential involvement in the regulation of hippocampal synaptic plasticity is an exciting avenue given that phosphodiesterase inhibition is a promising strategy to enhance synaptic plasticity and memory (Baillie et al. 2019; Barad et al. 1998). Further, previous studies have implicated PDE4A5 in mediating the impact of sleep deprivation on hippocampal synaptic plasticity and memory (Vecsey et al. 2009; Wong et al. 2019) and it therefore will be interesting to investigate whether POPDC1 has any role in sleep-related synaptic plasticity. We are in the process of generating conditional knockouts of *Popdc1* and these would be invaluable in investigating the function of POPDC1 protein in specific brain regions involved in learning and memory.

## Acknowledgements

We thank Ursula Herbort-Brand for the excellent technical assistance. We also thank Tania Chatterjee Chowdhury, Jia Ern Ong and Achala Thippeswamy for their help with colony maintenance and genotyping.

## Author Contributions

M.S.S and L.R performed the electrophysiology experiments. M.S.S performed the cAMP assays, analyzed the data and wrote the manuscript with inputs from other authors. T.A supervised electrophysiology experimental design, data analysis and provided inputs to the manuscript. R.F.R.S performed the Western blot analysis and L.F performed the LacZ immunohistochemistry. L.R, K.P.G and K.M performed some of the initial analysis of hippocampus function and memory in *Popdc1* KO mutants. T.B initiated this study and was involved in data interpretation and manuscript preparation.

## References

Abel, T., Martin, K. C., Bartsch, D., & Kandel, E. R. (1998). Memory suppressor genes: inhibitory constraints on the storage of long-term memory. Science, 279(5349), 338–341. doi:10.1126/science.279.5349.338

Abel, T., & Nguyen, P. V. (2008). Regulation of hippocampus-dependent memory by cyclic AMP-dependent protein kinase. Prog Brain Res, 169, 97–115. doi:10.1016/S0079-6123(07)00006-4

Abel, T., Nguyen, P. V., Barad, M., Deuel, T. A. S., Kandel, E. R., & Bourtchouladze, R. (1997). Genetic Demonstration of a Role for PKA in the Late Phase of LTP and in Hippocampus-Based Long-Term Memory. 88(5), 615–626. doi:10.1016/s0092-8674(00)81904-2

Andrée, B., Fleige, A., Arnold, H. H., & Brand, T. (2002). Mouse Pop1 is required for muscle regeneration in adult skeletal muscle. Mol Cell Biol, 22(5), 1504–1512.

Andree, B., Hillemann, T., Kessler-Icekson, G., Schmitt-John, T., Jockusch, H., Arnold, H. H., & Brand, T. (2000). Isolation and characterization of the novel popeye gene family expressed in skeletal muscle and heart. Dev Biol, 223(2), 371–382. doi:10.1006/dbio.2000.9751

Bach, M. E., Barad, M., Son, H., Zhuo, M., Lu, Y. F., Shih, R., … Kandel, E. R. (1999). Age-related defects in spatial memory are correlated with defects in the late phase of hippocampal long-term potentiation in vitro and are attenuated by drugs that enhance the cAMP signaling pathway. Proc Natl Acad Sci U S A, 96(9), 5280–5285. doi:10.1073/pnas.96.9.5280

Baillie, G. S., Sood, A., McPhee, I., Gall, I., Perry, S. J., Lefkowitz, R. J., & Houslay, M. D. (2003). beta-Arrestin-mediated PDE4 cAMP phosphodiesterase recruitment regulates beta-adrenoceptor switching from Gs to Gi. Proc Natl Acad Sci U S A, 100(3), 940–945. doi:10.1073/pnas.262787199

Baillie, G. S., Tejeda, G. S., & Kelly, M. P. (2019). Therapeutic targeting of 3’,5’-cyclic nucleotide phosphodiesterases: inhibition and beyond. Nat Rev Drug Discov, 18(10), 770–796. doi:10.1038/s41573-019-0033-4

Barad, M., Bourtchouladze, R., Winder, D. G., Golan, H., & Kandel, E. (1998). Rolipram, a type IV-specific phosphodiesterase inhibitor, facilitates the establishment of long-lasting long-term potentiation and improves memory. Proc Natl Acad Sci U S A, 95(25), 15020–15025. doi:10.1073/pnas.95.25.15020

Bauman, A. L., Goehring, A. S., & Scott, J. D. (2004). Orchestration of synaptic plasticity through AKAP signaling complexes. Neuropharmacology, 46(3), 299–310. doi:10.1016/j.neuropharm.2003.09.016

Benesh, E. C., Miller, P. M., Pfaltzgraff, E. R., Grega-Larson, N. E., Hager, H. A., Sung, B. H., … Bader, D. M. (2013). Bves and NDRG4 regulate directional epicardial cell migration through autocrine extracellular matrix deposition. Mol Biol Cell, 24(22), 3496–3510. doi:10.1091/mbc.E12-07-0539

Billard, J. M. (2010). Long-term depression in the hippocampal CA1 area of aged rats, revisited: contribution of temporal constraints related to slice preparation. PLoS One, 5(3), e9843. doi:10.1371/journal.pone.0009843

Blitzer, R. D., Wong, T., Nouranifar, R., Iyengar, R., & Landau, E. M. (1995). Postsynaptic cAMP pathway gates early LTP in hippocampal CA1 region. Neuron, 15(6), 1403–1414. doi:10.1016/0896-6273(95)90018-7

Bolger, G. B., Dunlop, A. J., Meng, D., Day, J. P., Klussmann, E., Baillie, G. S., … Houslay, M. D. (2015). Dimerization of cAMP phosphodiesterase-4 (PDE4) in living cells requires interfaces located in both the UCR1 and catalytic unit domains. Cell Signal, 27(4), 756–769. doi:10.1016/j.cellsig.2014.12.009

Brand, T. (2018). The Popeye Domain Containing Genes and Their Function as cAMP Effector Proteins in Striated Muscle. J Cardiovasc Dev Dis, 5(1). doi:10.3390/jcdd5010018

Buzsaki, G. (2005). Theta rhythm of navigation: link between path integration and landmark navigation, episodic and semantic memory. Hippocampus, 15(7), 827–840. doi:10.1002/hipo.20113

Cai, Y., Peng, Z., Guo, H., Wang, F., & Zeng, Y. (2017). TREK-1 pathway mediates isoflurane-induced memory impairment in middle-aged mice. Neurobiol Learn Mem, 145, 199–204. doi:10.1016/j.nlm.2017.10.012

Cedervall, P., Aulabaugh, A., Geoghegan, K. F., McLellan, T. J., & Pandit, J. (2015). Engineered stabilization and structural analysis of the autoinhibited conformation of PDE4. Proc Natl Acad Sci U S A, 112(12), E1414–1422. doi:10.1073/pnas.1419906112

Chavez-Noriega, L. E., & Stevens, C. F. (1992). Modulation of synaptic efficacy in field CA1 of the rat hippocampus by forskolin. Brain Res, 574(1-2), 85–92. doi:10.1016/0006-8993(92)90803-h

Chavez-Noriega, L. E., & Stevens, C. F. (1994). Increased transmitter release at excitatory synapses produced by direct activation of adenylate cyclase in rat hippocampal slices. J Neurosci, 14(1), 310–317. doi:10.1523/JNEUROSCI.14-01-00310.1994

Chen, A., Muzzio, I. A., Malleret, G., Bartsch, D., Verbitsky, M., Pavlidis, P., … Kandel, E. R. (2003). Inducible enhancement of memory storage and synaptic plasticity in transgenic mice expressing an inhibitor of ATF4 (CREB-2) and C/EBP proteins. Neuron, 39(4), 655–669. doi:10.1016/s0896-6273(03)00501-4

De Ridder, W., Nelson, I., Asselbergh, B., De Paepe, B., Beuvin, M., Ben Yaou, R., … Baets, J. (2019). Muscular dystrophy with arrhythmia caused by loss-of-function mutations in BVES. Neurol Genet, 5(2), e321. doi:10.1212/NXG.0000000000000321

Delgado, J. Y., Fink, A. E., Grant, S. G. N., O’Dell, T. J., & Opazo, P. (2018). Rapid homeostatic downregulation of LTP by extrasynaptic GluN2B receptors. J Neurophysiol, 120(5), 2351–2357. doi:10.1152/jn.00421.2018

Dudek, S. M., & Bear, M. F. (1992). Homosynaptic long-term depression in area CA1 of hippocampus and effects of N-methyl-D-aspartate receptor blockade. Proc Natl Acad Sci U S A, 89(10), 4363–4367. doi:10.1073/pnas.89.10.4363

Engmann, O., Campbell, J., Ward, M., Giese, K. P., & Thompson, A. J. (2010). Comparison of a protein-level and peptide-level labeling strategy for quantitative proteomics of synaptosomes using isobaric tags. J Proteome Res, 9(5), 2725–2733. doi:10.1021/pr900627e

Enyedi, P., & Czirjak, G. (2010). Molecular background of leak K+ currents: two-pore domain potassium channels. Physiol Rev, 90(2), 559–605. doi:10.1152/physrev.00029.2009

Frey, U., Huang, Y. Y., & Kandel, E. R. (1993). Effects of cAMP simulate a late stage of LTP in hippocampal CA1 neurons. Science, 260(5114), 1661–1664. doi:10.1126/science.8389057

Froese, A., Breher, S. S., Waldeyer, C., Schindler, R. F., Nikolaev, V. O., Rinne, S., … Brand, T. (2012). Popeye domain containing proteins are essential for stress-mediated modulation of cardiac pacemaking in mice. J Clin Invest, 122(3), 1119–1130. doi:10.1172/JCI59410

Hager, H. A., & Bader, D. M. (2009). Bves: ten years after. Histol Histopathol, 24(6), 777–787.

Hager, H. A., Roberts, R. J., Cross, E. E., Proux-Gillardeaux, V., & Bader, D. M. (2010). Identification of a novel Bves function: regulation of vesicular transport. EMBO J, 29(3), 532–545. doi:10.1038/emboj.2009.379

Havekes, R., Park, A. J., Tolentino, R. E., Bruinenberg, V. M., Tudor, J. C., Lee, Y., … Abel, T. (2016). Compartmentalized PDE4A5 Signaling Impairs Hippocampal Synaptic Plasticity and Long-Term Memory. J Neurosci, 36(34), 8936–8946. doi:10.1523/JNEUROSCI.0248-16.2016

Heynen, A. J., Abraham, W. C., & Bear, M. F. (1996). Bidirectional modification of CA1 synapses in the adult hippocampus in vivo. Nature, 381(6578), 163–166. doi:10.1038/381163a0

Honore, E. (2007). The neuronal background K2P channels: focus on TREK1. Nat Rev Neurosci, 8(4), 251–261. doi:10.1038/nrn2117

Houslay, M. D. (2010). Underpinning compartmentalised cAMP signalling through targeted cAMP breakdown. Trends Biochem Sci, 35(2), 91–100. doi:10.1016/j.tibs.2009.09.007

Huang, Y. Y., & Kandel, E. R. (1994). Recruitment of long-lasting and protein kinase A-dependent long-term potentiation in the CA1 region of hippocampus requires repeated tetanization. Learn Mem, 1(1), 74–82.

Indrawati, L. A., Iida, A., Tanaka, Y., Honma, Y., Mizoguchi, K., Yamaguchi, T., … Nishino, I. (2020). Two Japanese LGMDR25 patients with a biallelic recurrent nonsense variant of BVES. Neuromuscul Disord, 30(8), 674–679. doi:10.1016/j.nmd.2020.06.004

Jackman, S. L., & Regehr, W. G. (2017). The Mechanisms and Functions of Synaptic Facilitation. Neuron, 94(3), 447–464. doi:10.1016/j.neuron.2017.02.047

Kelly, M. P. (2018). Cyclic nucleotide signaling changes associated with normal aging and age-related diseases of the brain. Cell Signal, 42, 281–291. doi:10.1016/j.cellsig.2017.11.004

Kramar, E. A., Chen, L. Y., Brandon, N. J., Rex, C. S., Liu, F., Gall, C. M., & Lynch, G. (2009). Cytoskeletal changes underlie estrogen’s acute effects on synaptic transmission and plasticity. J Neurosci, 29(41), 12982–12993. doi:10.1523/JNEUROSCI.3059-09.2009

Malleret, G., Haditsch, U., Genoux, D., Jones, M. W., Bliss, T. V., Vanhoose, A. M., … Mansuy, I. M. (2001). Inducible and reversible enhancement of learning, memory, and long-term potentiation by genetic inhibition of calcineurin. Cell, 104(5), 675–686. doi:10.1016/s0092-8674(01)00264-1

McPhee, I., Cochran, S., & Houslay, M. D. (2001). The novel long PDE4A10 cyclic AMP phosphodiesterase shows a pattern of expression within brain that is distinct from the long PDE4A5 and short PDE4A1 isoforms. Cell Signal, 13(12), 911–918. doi:10.1016/s0898-6568(01)00217-0

Neves, G., Cooke, S. F., & Bliss, T. V. (2008). Synaptic plasticity, memory and the hippocampus: a neural network approach to causality. Nat Rev Neurosci, 9(1), 65–75. doi:10.1038/nrn2303

Nguyen, P. V., Abel, T., & Kandel, E. R. (1994). Requirement of a critical period of transcription for induction of a late phase of LTP. Science, 265(5175), 1104–1107. doi:10.1126/science.8066450

Nguyen, P. V., & Kandel, E. R. (1997). Brief theta-burst stimulation induces a transcription-dependent late phase of LTP requiring cAMP in area CA1 of the mouse hippocampus. Learn Mem, 4(2), 230–243. doi:10.1101/lm.4.2.230

Nguyen, P. V., & Woo, N. H. (2003). Regulation of hippocampal synaptic plasticity by cyclic AMP-dependent protein kinases. Progress in Neurobiology, 71(6), 401–437. doi:10.1016/j.pneurobio.2003.12.003

Nie, T., McDonough, C. B., Huang, T., Nguyen, P. V., & Abel, T. (2007). Genetic disruption of protein kinase A anchoring reveals a role for compartmentalized kinase signaling in theta-burst long-term potentiation and spatial memory. J Neurosci, 27(38), 10278–10288. doi:10.1523/JNEUROSCI.1602-07.2007

Osler, M. E., Chang, M. S., & Bader, D. M. (2005). Bves modulates epithelial integrity through an interaction at the tight junction. J Cell Sci, 118(Pt 20), 4667–4678.

Otmakhov, N., Khibnik, L., Otmakhova, N., Carpenter, S., Riahi, S., Asrican, B., & Lisman, J. (2004). Forskolin-induced LTP in the CA1 hippocampal region is NMDA receptor dependent. J Neurophysiol, 91(5), 1955–1962. doi:10.1152/jn.00941.2003

Patriarchi, T., Buonarati, O. R., & Hell, J. W. (2018). Postsynaptic localization and regulation of AMPA receptors and Cav1.2 by beta2 adrenergic receptor/PKA and Ca(2+)/CaMKII signaling. EMBO J, 37(20), e99771. doi:10.15252/embj.201899771

Perry, S. J., Baillie, G. S., Kohout, T. A., McPhee, I., Magiera, M. M., Ang, K. L., … Lefkowitz, R. J. (2002). Targeting of cyclic AMP degradation to beta 2-adrenergic receptors by beta-arrestins. Science, 298(5594), 834–836. doi:10.1126/science.1074683

Regehr, W. G. (2012). Short-term presynaptic plasticity. Cold Spring Harb Perspect Biol, 4(7), a005702. doi:10.1101/cshperspect.a005702

Rinne, S., Ortiz-Bonnin, B., Stallmeyer, B., Kiper, A. K., Fortmuller, L., Schindler, R. F. R., … Decher, N. (2020). POPDC2 a novel susceptibility gene for conduction disorders. J Mol Cell Cardiol, 145, 74–83. doi:10.1016/j.yjmcc.2020.06.005

Schindler, R. F., & Brand, T. (2016). The Popeye domain containing protein family - A novel class of cAMP effectors with important functions in multiple tissues. Prog Biophys Mol Biol, 120(1-3), 28–36. doi:10.1016/j.pbiomolbio.2016.01.001

Schindler, R. F., Scotton, C., Zhang, J., Passarelli, C., Ortiz-Bonnin, B., Simrick, S., … Ferlini, A. (2016). POPDC1(S201F) causes muscular dystrophy and arrhythmia by affecting protein trafficking. J Clin Invest, 126(1), 239–253. doi:10.1172/JCI79562

Schleicher, K., & Zaccolo, M. (2018). Using cAMP Sensors to Study Cardiac Nanodomains. J Cardiovasc Dev Dis, 5(1). doi:10.3390/jcdd5010017

Schleicher, K., & Zaccolo, M. (2020). Axelrod Symposium 2019: Phosphoproteomic analysis of G-protein-coupled pathways. Mol Pharmacol, 99(5), 383–391. doi:10.1124/mol.119.118869

Shetty, M. S., Sharma, M., Hui, N. S., Dasgupta, A., Gopinadhan, S., & Sajikumar, S. (2015). Investigation of Synaptic Tagging/Capture and Cross-capture using Acute Hippocampal Slices from Rodents. J Vis Exp, 103, e53008. doi:10.3791/53008

Staubli, U. V., & Ji, Z. X. (1996). The induction of homo- vs. heterosynaptic LTD in area CA1 of hippocampal slices from adult rats. Brain Res, 714(1-2), 169–176. doi:10.1016/0006-8993(95)01523-x

Swan, A. H., Gruscheski, L., Boland, L. A., & Brand, T. (2019). The Popeye domain containing gene family encoding a family of cAMP-effector proteins with important functions in striated muscle and beyond. J Muscle Res Cell Motil. doi:10.1007/s10974-019-09523-z

Tibbo, A. J., Dobi, S., McFall, A., Tejeda, G. S., Blair, C., MacLeod, R., … Baillie, G. S. (2020). Phosphodiesterase Type 4 anchoring regulates cAMP signaling to Popeye domain-containing proteins. bioRxiv, 2020.2009.2010.290825. doi:10.1101/2020.09.10.290825

Vasavada, T. K., DiAngelo, J. R., & Duncan, M. K. (2004). Developmental expression of Pop1/Bves. J Histochem Cytochem, 52(3), 371–377.

Vecsey, C. G., Baillie, G. S., Jaganath, D., Havekes, R., Daniels, A., Wimmer, M., … Abel, T. (2009). Sleep deprivation impairs cAMP signalling in the hippocampus. Nature, 461(7267), 1122–1125. doi:10.1038/nature08488

Vissing, J., Johnson, K., Topf, A., Nafissi, S., Diaz-Manera, J., French, V. M., … Straub, V. (2019). POPDC3 Gene Variants Associate with a New Form of Limb Girdle Muscular Dystrophy. Ann Neurol, 86(6), 832–843. doi:10.1002/ana.25620

Wang, W., Kiyoshi, C. M., Du, Y., Taylor, A. T., Sheehan, E. R., Wu, X., & Zhou, M. (2020). TREK-1 Null Impairs Neuronal Excitability, Synaptic Plasticity, and Cognitive Function. Mol Neurobiol, 57(3), 1332–1346. doi:10.1007/s12035-019-01828-x

Weng, W., Chen, Y., Wang, M., Zhuang, Y., & Behnisch, T. (2016). Potentiation of Schaffer-Collateral CA1 Synaptic Transmission by eEF2K and p38 MAPK Mediated Mechanisms. Front Cell Neurosci, 10, 247. doi:10.3389/fncel.2016.00247

Wong, L. W., Tann, J. Y., Ibanez, C. F., & Sajikumar, S. (2019). The p75 Neurotrophin Receptor Is an Essential Mediator of Impairments in Hippocampal-Dependent Associative Plasticity and Memory Induced by Sleep Deprivation. J Neurosci, 39(28), 5452–5465. doi:10.1523/JNEUROSCI.2876-18.2019

Zaccolo, M., Zerio, A., & Lobo, M. J. (2021). Subcellular Organization of the cAMP Signaling Pathway. Pharmacol Rev, 73(1), 278–309. doi:10.1124/pharmrev.120.000086

Zucker, R. S., & Regehr, W. G. (2002). Short-term synaptic plasticity. Annu Rev Physiol, 64, 355–405. doi:10.1146/annurev.physiol.64.092501.114547

